# Geometric confinement regulates ERK signaling dynamics and collective migration

**DOI:** 10.64898/2026.06.04.730094

**Authors:** Marie Versaevel, Rémi Tranzer, Marine Luciano, Edouard Hannezo, Tsuyoshi Hirashima, Sylvain Gabriele

## Abstract

Collective cell migration is a fundamental process in morphogenesis, tissue repair, cancer invasion, and frequently occurs under geometric confinement in vivo. However, how confinement interfaces with signaling pathways that coordinate collective motion remains poorly understood. Here, we confine migrating epithelial monolayers within adhesive microstripes of defined width and observe a progressive slow-down of collective migration with increasing spatial confinement. Combining biophysical modeling, live imaging of ERK activity, and pharmacological perturbations, we show that confinement increases tissue crowding while reducing cell and nuclear projected areas, thereby shifting epithelial tissues toward a mechanically compressed state associated with dampened ERK waves. Across conditions, migration speed scales with ERK signaling dynamics, which correlates with EGFR signaling as well as cell and nuclear projected areas, together serving as quantitative proxies for the confinement-imposed mechanical state. Pharmacological inhibition of ROCK restores cell spreading, ERK signaling, and migration under strong confinement, demonstrating that this state is reversible and governed by actomyosin contractility. Together, our results identify geometric confinement as a physical regulator of a contractility-dependent mechanochemical state that controls ERK signaling and collective migration in epithelial tissues.

---

Collective cell migration is a fundamental process underlying tissue morphogenesis, wound healing, and cancer invasion. In physiological contexts, migrating epithelial tissues rarely move in unbounded environments but instead navigate through geometrically constrained spaces defined by surrounding tissues and extracellular matrix architecture. Increasing evidence indicates that confinement is not merely a passive boundary condition but an active regulator of collective dynamics, influencing cell coordination, force transmission, polarity, and tissue organization (1–8). Recent work further shows that geometric confinement and tissue crowding can profoundly alter epithelial physical states, including cell density, cell shape, mechanical stress distribution, and transitions between fluid-like and jammed migratory regimes (9–11). However, how confinement-induced changes in epithelial physical state quantitatively couple to ERK signaling dynamics that regulates collective migration remains poorly understood.

The ERK/MAPK cascade is a central regulator of cell behavior, controlling proliferation, survival, differentiation, and migration in response to extracellular cues (12). In epithelial collectives, ERK activity propagates as spatiotemporal waves that coordinate cell motion, polarity, survival, and tissue homeostasis (13–16). These signaling dynamics are part of mechanochemical feedback loops in which mechanical forces activate RTK–ERK signaling, while ERK activity in turn regulates cytoskeletal forces and collective tissue motion (12, 17, 18).

Cell–cell and cell–matrix adhesions provide potential interfaces for this mechanochemical coupling. Adherens junctions not only maintain epithelial cohesion but also regulate collective polarity and migration, while E-cadherin–dependent mechanisms can control EGFR activation at cell–cell contacts (5, 19). In parallel, focal adhesions and nuclear mechanotransduction have recently been implicated in density-dependent ERK activation and epithelial migration, suggesting that epithelial tissues can convert changes in density and mechanical state into ERK-dependent migratory responses (20). Together, these studies suggest that ERK signaling is sensitive to the mechanical and adhesive state of epithelial tissues, yet how lateral confinement quantitatively tunes ERK activity during collective migration remains unclear.

Here, we address this question using adhesive microstripes of controlled width to systematically modulate epithelial tissue confinement over a broad range of spatial scales, from large multicellular tissues to dimensions approaching the width of a single cell. By combining live-cell imaging of ERK activity with quantitative analysis of collective migration, cell morphology, EGFR activation, and nuclear geometry, we show that geometric confinement induces a progressive reduction in ERK activity amplitude that strongly correlates with migration efficiency, while leaving the temporal characteristics of ERK dynamics largely unchanged. We further show that this modulation is associated with confinement-dependent changes in cell spreading, junctional EGFR signaling, and nuclear projected area, and that it can be reversed by inhibition of ROCK-dependent contractility. Together, our results identify geometric confinement as a physical regulator of a contractility-dependent mechanochemical state that governs ERK signaling and collective epithelial migration.

## Results

### Geometric confinement defines distinct regimes of collective epithelial migration

To determine how lateral confinement regulates collective epithelial migration, we developed a micropatterning assay in which confluent MDCK monolayers migrate from a large reservoir into fibronectin-coated adhesive microstripes of defined widths ranging from 500 to 15 µm **(Fig. 1a**,**b)**. This strategy builds on previous confinement-based migration assays showing that physical boundaries can organize collective epithelial motion (3), while extending this framework to stronger confinement regimes and to the subsequent analysis of ERK signaling dynamics. Before migration initiation, the fibronectin-coated microstripes were physically masked with a passivated PDMS sheet during cell seeding, thereby restricting epithelial cell adhesion and growth to the large reservoir region. Upon removal of the PDMS sheet, the adhesive stripes became exposed and synchronously triggered collective migration into the confined tracks. MDCK monolayers then progressively invaded the microstripes while remaining laterally constrained by the non-adhesive surrounding regions **(Fig. 1a)**, enabling collective migration to be monitored under well-controlled confinement widths ranging from 500 to 15 µm. n wide stripes, epithelial monolayers displayed rapid and persistent front progression, whereas collective migration progressively slowed as confinement increased and stripe width decreased **(Fig. 1b–d and SI Movie S1)**.

**Figure 1.**
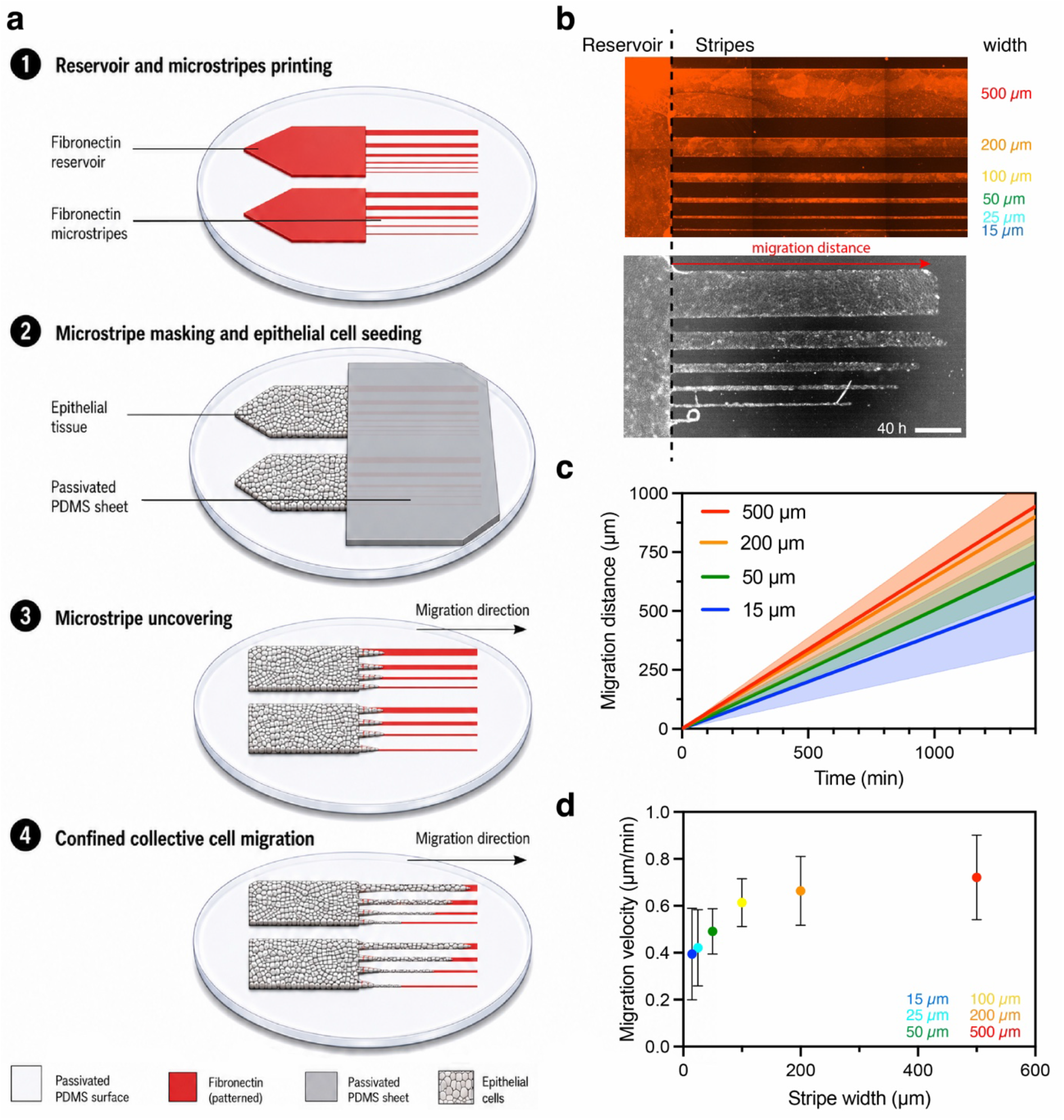
Geometric confinement defines distinct regimes of collective migration. **(a)** Experimental sketch of fibronectin (FN) microstripes of defined widths (500–15 µm) micropatterned on a PDMS-coated glass substrate and temporarily masked with a passivated PDMS sheet. MDCK cells are seeded within an FN-coated reservoir, and the removal of a PDMS sheet exposed the microstripes to initiate confined collective migration. **(b)** Representative images of (top) fluorescent labeled FN microstripes of varying widths connected to a reservoir (red) and (bottom) differential interference contrast (DIC) image of MDCK cells migrating along microstripes of decreasing widths (500-15 µm) after 40 h, illustrating confinement-dependent collective migration. Scale bar, 500 µm. **(c)** Migration distance as a function of time for four representative microstripe widths (500 µm, red; 200 µm, orange; 50 µm, green; and 15 µm, blue). Solid lines represent linear regressions of the mean ± SD from multiple independent experiments. **(d)** Mean migration velocity as a function of stripe width, revealing reduced collective migration under strong confinement. For (c) and (d), data were obtained from N=3 replicates, with n=9 microstripes (500 µm), n=13 (200 µm), n=10 (100 µm), n=10 (50 µm), n=8 (25 µm), and n=5 (15 µm).

Quantification of front displacement over time showed an approximately linear progression for each width **(SI Appendix, Fig. S1)**, allowing migration velocity to be extracted from the slope of the displacement curves **(Fig. 1c)**. Our results indicated that mean migration velocity depended strongly on stripe width **(Fig. 1d)**. Indeed, compared with wide tracks, strongly confined stripes supported markedly reduced migration, with the lowest velocities observed in the narrowest conditions. Particle image velocimetry (PIV) analysis further showed that velocity vectors remain predominantly aligned along the stripe axis across all widths **(SI Appendix, Fig. S2a–f and SI Movie S2)**, with confinement primarily affecting the magnitude rather than the direction of migration **(SI Appendix, Fig. S2g**,**h)**. Our data reveal that strong lateral confinement can instead impose a low-motility regime.

Together, these results establish spatial confinement as a key physical control parameter of collective migration, ranging from efficient migration in wide stripes to impaired collective advance under strong confinement. We next asked whether this confinement-dependent reduction in migration is specific to linear geometries or reflects a more general physical constraint.

### Confinement-dependent slowing of collective migration is geometry-independent

To determine whether the confinement-dependent reduction in migration observed in linear stripes is specific to this geometry or reflects a more general physical constraint, we next examined epithelial migration in funnel-shaped micropatterns, in which cells progressively encounter decreasing widths **(Fig. 2a)**. Time-lapse imaging revealed that MDCK monolayers collectively invaded the funnel structures, with cells advancing from wide regions toward increasingly confined zones **(SI Movie S3)**. As migration proceeded, the advancing front progressively slowed down as it entered narrower regions, despite continuous cell supply from the reservoir. This spatial deceleration was evident from the gradual accumulation of cells and reduced front progression in highly confined regions **(Fig. 2a)**. To directly compare migration dynamics across geometries, we quantified migration velocity of the leading front as a function of local confinement width in both uniform stripes and funnel patterns. Remarkably, velocities measured in funnels closely overlapped with those obtained in straight stripes of equivalent width **(Fig. 2b)**. In both cases, migration speed decreased monotonically with decreasing width, indicating that the effect of confinement is independent of the specific pattern geometry. Averaging across independent experiments confirmed a robust and reproducible relationship between confinement width and migration velocity **(Fig. 2c)**, demonstrating that local geometric constraints, rather than global pattern geometry, govern collective migration efficiency. Together, these findings demonstrate that confinement-dependent slowing of collective migration is a robust property governed by local physical constraints.

**Figure 2.**
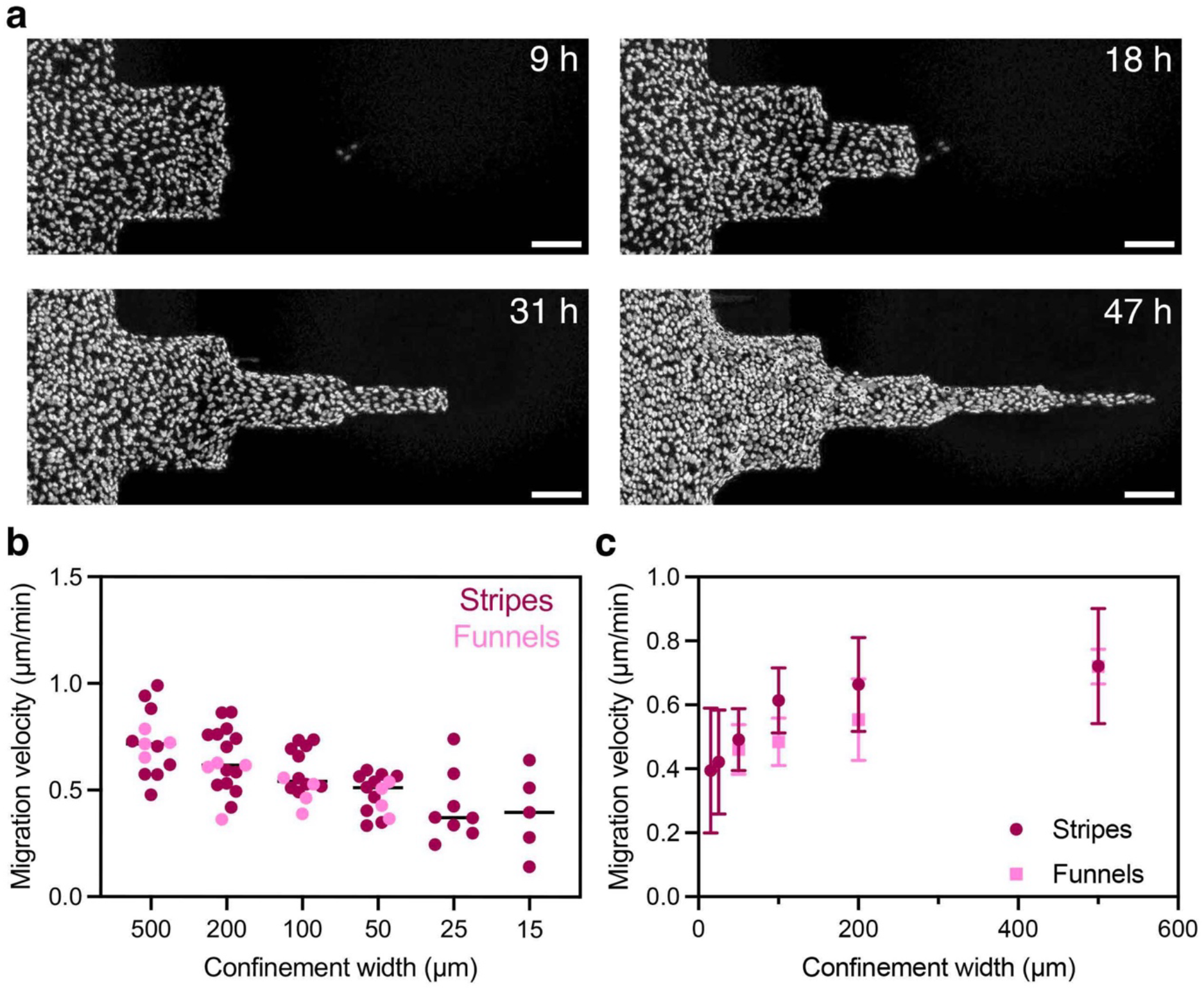
Confinement-dependent reduction of collective migration is robust across pattern geometries. (a) Time-lapse images of nuclei in MDCK monolayers migrating through funnel-shaped micropatterns at the indicated time points (9, 18, 31, and 47 h), illustrating progressive invasion into regions of increasing confinement corresponding to decreasing widths (500, 200, 100, 50, 25, and 15 µm). Scale bars, 200 µm. (b) Migration velocity as a function of local confinement width for cells migrating either along uniform microstripes (dark purple) or through funnel-shaped geometries (light purple). Individual measurements reveal a consistent reduction in migration velocity with increasing confinement across both micropattern designs. (c) Mean migration velocity as a function of confinement width for uniform microstripes (dark purple) and funnel-shaped geometries (light purple), demonstrating that the confinement-dependent reduction in collective migration is preserved across distinct micropattern architectures. Migration velocities in funnel geometries were obtained from n=4 independent funnel patterns across N=2 replicates.

### Cell and nuclear geometry serve as quantitative proxies for the confinement-imposed mechanical state

Having established that spatial confinement modulates cell migration speed at the collective level, we next asked if the regulation of migration speed is reflected in morphological changes, for example at the nuclear level, a key integrator of mechanical constraints at the cellular scale (21, 22).

Imaging of nuclei in migrating MDCK monolayers revealed pronounced confinement-dependent changes in nuclear architecture and orientation **(Fig. 3a)**. As stripe width decreased, nuclei became progressively aligned along the migration axis **(Fig. 3b)** and exhibited increased deformation, as quantified by the nuclear aspect ratio **(Fig. 3c, SI Movies S4-S5)**. Spatial confinement within tissues is also accompanied by significant changes in cell morphology (**SI Appendix, Fig. S3A)**. Cells in highly confined tissues adopt an elongated shape, whereas they appear more rounded in wider microstripes (**SI Appendix, Fig. S3B)**. Interestingly, nuclear deformation closely followed cell shape changes, with a good correlation between nuclear aspect ratio and cell shape index, CSI **(Fig. 3d**, R^2^ = 0.6594), indicating a tight mechanical coupling between cellular and nuclear architecture. In parallel, nuclear density increased under stronger confinement **(Fig. 3e)**, consistent with enhanced cell crowding in narrow microstripes. Higher nuclear density was in turn associated with greater nuclear elongation **(Fig. 3f)**, suggesting that both lateral geometric constraints and crowding cooperatively contribute to nuclear mechanical adaptation.

**Figure 3.**
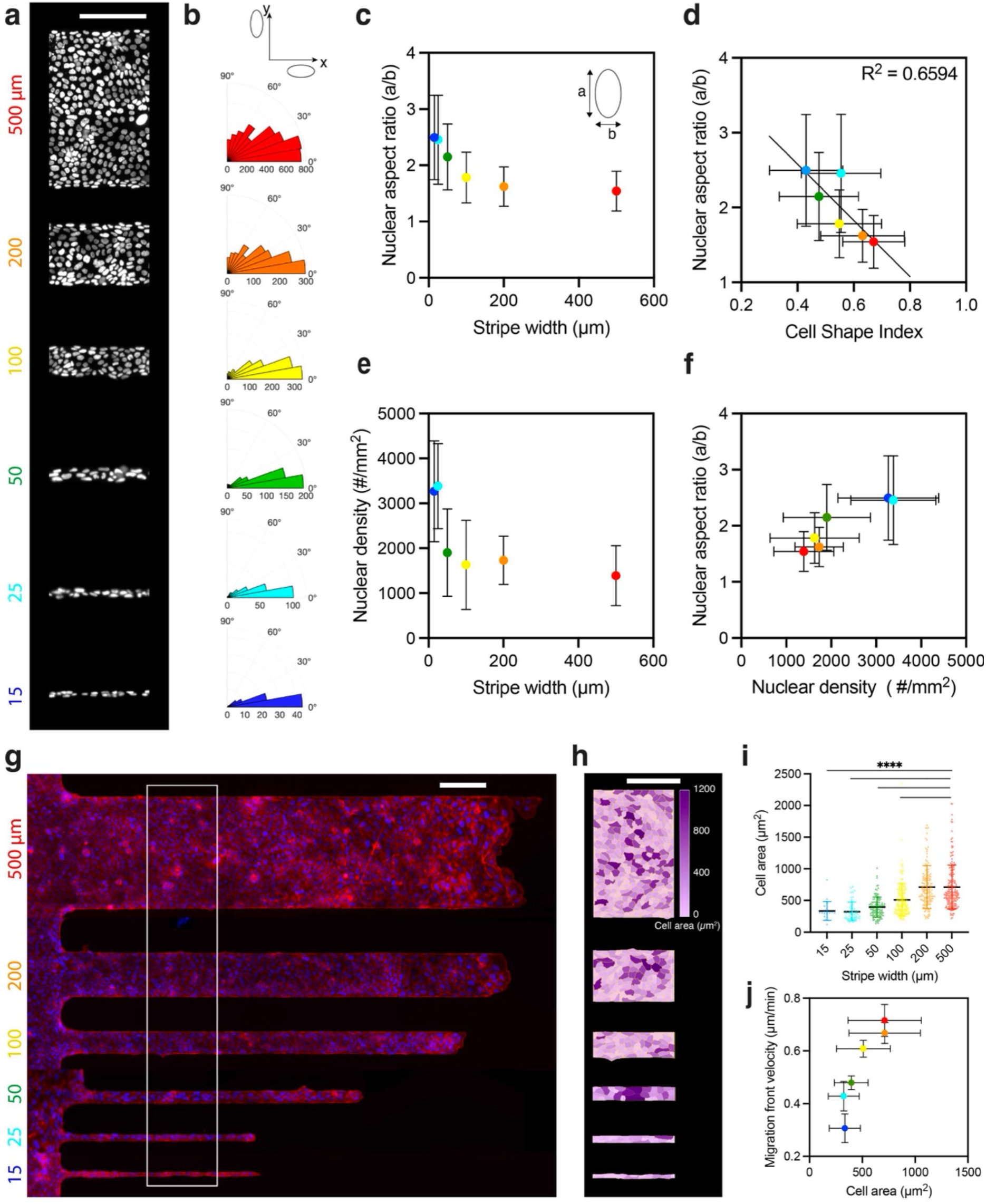
Spatial confinement progressively modulates nuclear organization and cell spreading. **(a)** Representative images of nuclei stained with Dapi in MDCK monolayers migrating along fibronectin microstripes of decreasing widths (500–15 µm), illustrating confinement-dependent changes in nuclear organization. Scale bar, 200 µm. **(b)** Angular distribution of nuclear orientation for each microstripe width, showing progressive alignment of nuclei along the microstripe axis under strong confinement. **(c)** Nuclear aspect ratio as a function of microstripe width, indicating progressive nuclear elongation with increasing confinement. **(d)** Nuclear aspect ratio as a function of cell shape index, revealing a strong coupling between nuclear and cellular geometry (R^2^ = 0.6594). **(e)** Nuclear density as a function of microstripe width, showing increased tissue crowding under strong confinement. Nuclear density measurements were obtained from n = 8 samples (500 µm), n = 5 (200 µm), n = 8 (100 µm), n = 7 (50 µm), n = 4 (25 µm), and n = 5 (15 µm). N ≥ 3 replicates for all conditions.**(f)** Nuclear aspect ratio as a function of nuclear density, indicating that increased crowding is associated with enhanced nuclear deformation. Nuclear aspect ratio were obtained from n = 2686 nuclei (500 µm), n = 21063 (200 µm), n = 715 (100 µm), n = 546 (50 µm), n = 118 (25 µm), and n = 49 (15 µm). N ≥ 3 replicates for all conditions. **(g)** Representative reconstructed fluorescence image of MDCK monolayers migrating along fibronectin microstripes of varying widths and immunostained for nuclei (blue, DAPI) and cell–cell junctions (red, β-catenin). The white box indicates the region used for cell area quantification. Scale bar, 200 µm. **(h)** Spatial maps of cell area across microstripes of decreasing widths, highlighting confinement-dependent reductions in cell spreading. Scale bar, 200 µm. **(i)** Quantification of cell area as a function of microstripe width, showing a progressive decrease in cell spreading with increasing confinement. Cell area measurements were obtained from 3 independent experiments, with n = 197 cells (500 µm), n = 155 (200 µm), n = 186 (100 µm), n = 111 (50 µm), n = 52 (25 µm), and n = 20 (15 µm). N ≥ 3 replicates for all conditions. **(j)** Migration front velocity as a function of mean cell area, revealing a positive correlation between collective migration efficiency and cell spreading. Points are color-coded according to microstripe width.

Given the important role of cell area in ERK signaling (23), we next examined the impact of the spatial confinement on cell morphology. MDCK monolayers migrating along fibronectin stripes of varying widths were fixed and stained for cell–cell junctions **(Fig. 3g)**, enabling quantitative analysis of cell shape and spreading **(Fig. 3h)**. Spatial mapping revealed pronounced confinement-dependent changes in cell morphology, with cells progressively reducing their spreading area as stripe width decreased **(Fig. 3i and SI Appendix Figs. 3c)**, indicating that lateral geometric constraints restrict cell spreading within migrating epithelial tissues. Consistently, nuclear projected area progressively decreased with increasing confinement **(SI Appendix Figs. 3d-e)**, revealing the strong sensitivity of nuclear organization to spatial constraints. In addition to nuclear elongation and alignment along the microstripe axis, confinement therefore induces a marked reduction in nuclear projected area.

Finally, our results reveal that migration speed positively correlates with cell spreading area **(Fig. 3j)**, suggesting that confinement-induced changes in cellular and nuclear mechanical state may regulate mechanosensitive signaling pathways controlling collective migration. Together, these observations indicate that confinement progressively shifts epithelial tissues toward a mechanically crowded and weakly spread state. Because ERK signaling is known to respond to cell deformation and spreading, we next investigated how confinement modulates ERK dynamics during collective migration.

### Confinement progressively dampens ERK signaling and migration

To address this possibility, we monitored ERK activity in migrating MDCK monolayers stably expressing the EKARrEV-NLS FRET biosensor (24–26) over extended periods of time, up to 4 days **(Fig. 4a)**. This approach enabled quantitative mapping of ERK dynamics at single-cell resolution during collective migration **(Fig. 4b-c)**. Spatiotemporal analysis revealed that ERK activity amplitude was strongly dependent on confinement **(Fig. 4d and SI Appendix, Fig. 4a–d)**. In wide microstripes, pronounced ERK activation waves propagated collectively across the migrating epithelium **(SI Movie S6)**, whereas highly confined stripes displayed strongly dampened and disorganized ERK oscillatory dynamics, with only weak residual fluctuations remaining **(SI Movie S7)**.

**Figure 4.**
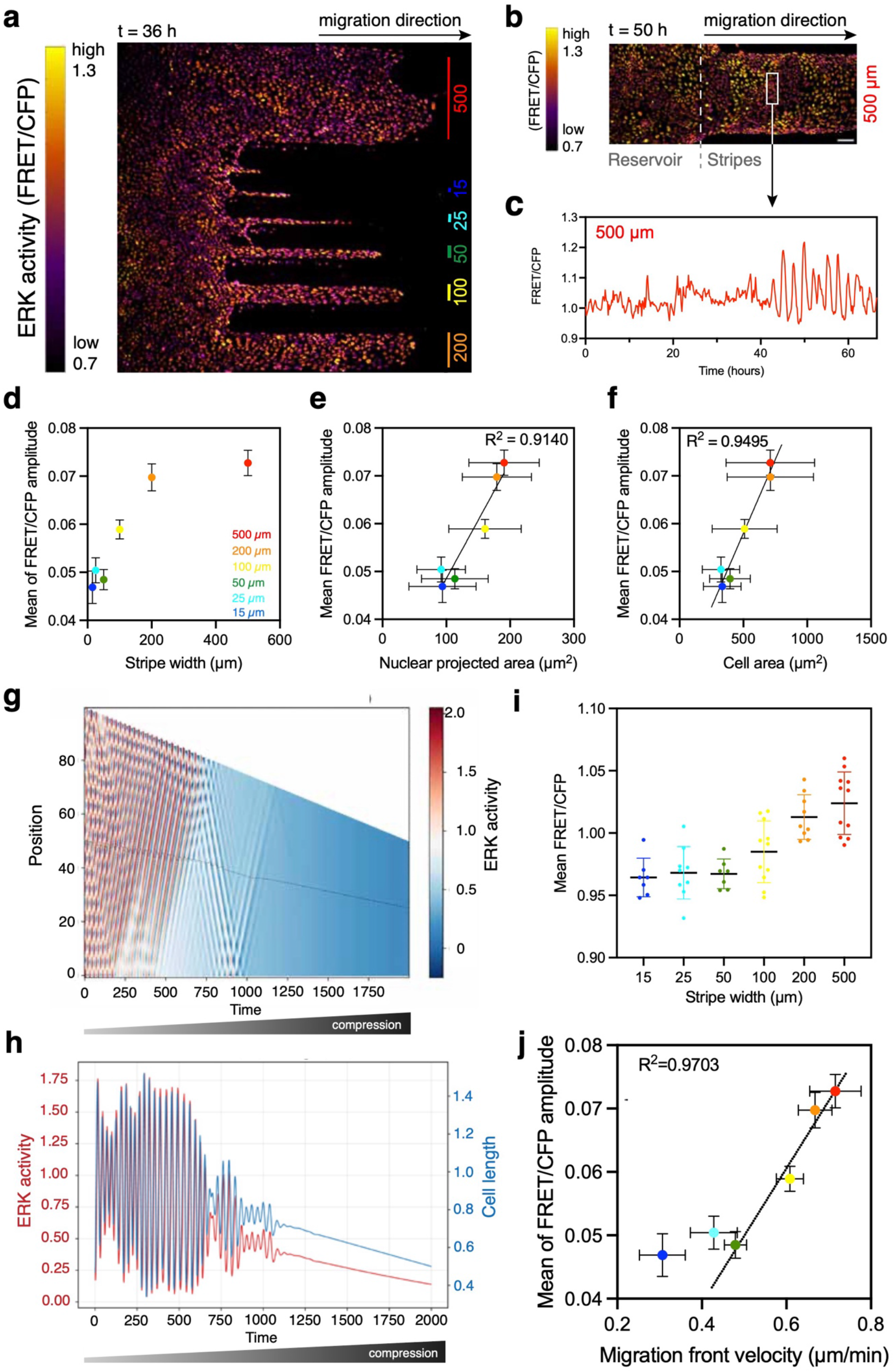
ERK activity is progressively reduced by confinement and scales with collective migration speed. **(a)** Representative color-coded map of ERK activity (FRET/CFP) at 36 h in MDCK monolayers migrating along fibronectin microstripes of decreasing widths, revealing confinement-dependent spatial patterns of ERK signaling. **(b)** Higher-magnification view of a wide microstripe (500 µm) at 50 h, with ERK activity mapped at single-cell resolution. The dashed line indicates the boundary between the reservoir and the microstripe, and the white box highlights a representative region of interest (ROI). The black arrow indicates the direction of migration. Scale bar, 100 µm. **(c)** Representative temporal trace of ERK activity (FRET/CFP) measured in the ROI shown in (b), revealing oscillatory ERK dynamics over time. **(d)** Mean ERK activity amplitude as a function of microstripe width, demonstrating a confinement-dependent reduction in ERK oscillation amplitude. **(e)** Mean ERK activity amplitude as a function of nuclear projected area, revealing a strong positive correlation (R^2^ = 0.9140), indicating that nuclear projected area provides a quantitative proxy for the confinement-imposed mechanical state. **(f)** Mean ERK activity amplitude as a function of cellular area, showing that increased cell spreading is associated with higher ERK activity amplitude (R^2^ = 0.9495). **(g-h)** Theoretical kymographs **(g)** generated from a mechanochemical model of ERK waves during progressively increasing confinement/compression. As confinement gradually increases over time, the model predicts a progressive reduction in ERK activity followed by suppression of oscillatory wave dynamics beyond a critical confinement threshold (see SI Theory). Colors represent ERK activity levels, with red indicating high ERK activity and blue indicating low ERK activity. Alternating diagonal bands correspond to propagating ERK waves. Panel h shows how ERK activity (red) and cellular length (blue) evolves over time for a single cell (followed by a dashed line in panel g). **(i)** Mean ERK activity (mean FRET/CFP ratio) as a function of microstripe width, revealing a progressive reduction in mean ERK activity under confinement, consistent with the theoretical prediction shown in (g). **(j)** Mean ERK activity amplitude as a function of migration front velocity, revealing a strong positive correlation (R^2^ = 0.9703) between ERK oscillatory amplitude and collective migration efficiency above 50 µm, with oscillations collapsing under strong confinement (< 50 µm). FRET/CFP signals were acquired from N replicates, with a total of n measurements: N = 9 and n = 373 (500 µm), N = 7 and n = 279 (200 µm), N = 9 and n = 411 (100 µm), N = 5 and n = 222 (50 µm), N = 6 and n = 171 (25 µm), and N = 4 and n = 109 (15 µm).

Because ERK signaling in migrating epithelial tissues is often organized as pulsatile waves, we quantified ERK activity amplitude from local maxima and minima of the FRET/CFP signal as a robust readout of signaling strength **(SI Appendix, Fig. 4a–c)**. Strikingly, ERK activity amplitude strongly correlated with nuclear projected area across all conditions (**Fig. 4e**; R^2^ = 0.9140), with larger projected nuclear areas associated with higher ERK signaling levels. Importantly, this reduction in projected area with confinement occurs without major changes in nuclear volume **(SI Appendix, Fig. S5f-g, SI Movies S4-S5**, indicating that nuclear shape remodeling is accommodated through three-dimensional reorganization and an increase in nuclear height **(SI Appendix, Fig. S5a-e)**.

The mean ERK activity amplitude decreased monotonically with increasing confinement **(Fig. 4e and SI Appendix, Figs. 4d-e)**, indicating that strong geometric constraints dampen ERK signaling. In contrast, the time between successive ERK activity maxima remained largely independent of stripe width **(SI Appendix, Fig. 4f)**, showing that confinement primarily modulates ERK amplitude rather than wave period.

Given that average ERK activity has previously been shown to be mechanosensitive and positively correlated with cell area in mechanically stretched epithelia(14), we next asked whether ERK signaling similarly scales with cell spreading under spatial confinement. Indeed, ERK activity amplitude strongly positively correlated with cell area across all confinement conditions **(Fig. 4f**, R^2^ = 0.9495), suggesting that cell spreading acts as a quantitative proxy of the confinement-imposed mechanical state governing ERK signaling dynamics.

Consistent with this idea, ERK activity amplitude also positively correlated with nuclear aspect ratio **(SI Appendix, Fig. S6a**, R^2^ = 0.8607) and showed a dependence on nuclear density **(SI Appendix, Fig. S6b)**, particularly at low-density regimes. Together, these observations indicate that ERK signaling scales with multiple geometric proxies of the confinement-imposed mechanical state of epithelial tissues.

To interpret this relationship mechanistically, we examined the effect of progressive confinement in a previously published mechanochemical model of ERK waves **(Figs. 4g-h)** (27). In this framework, increasing confinement progressively reduces the effective area available to each cell, thereby shifting the system toward a mechanically compressed state associated with lower average ERK activity **(Fig. 4g)**. Consistent with this theoretical prediction, experimental quantification revealed a progressive reduction in mean FRET/CFP levels with increasing confinement **(Fig. 4i)**. Importantly, the model further predicts that beyond a critical level of confinement, the oscillatory ERK regime becomes strongly dampened, while the intrinsic oscillation period remains largely preserved **(Fig. 4h)**. Mechanistically, this transition emerges because progressive compression shifts the system outside the oscillatory regime by saturating the mechanochemical feedback underlying ERK waves. Consistent with this theoretical prediction, ERK activity amplitude strongly correlated with migration front velocity across confinement conditions **(Fig. 4j**, R^2^=0.9703), with a marked reduction in oscillatory ERK activity observed under strong confinement.

Next, we explored within the same theoretical framework how suppression of ERK waves could contribute to the reduction in collective migration observed under confinement. Previous studies have suggested that ERK inhibition can locally increase traction forces, which could in principle favor migration (14). However, ERK waves were also proposed to provide directional cues required for coordinated collective motion. Incorporating these two effects into the previously calibrated mechanochemical model suggested that suppression of ERK waves under confinement would impair directional collective migration, even if individual cell traction forces locally increase **(see SI Theory)**. In this framework, the loss of spatiotemporal ERK coordination prevents efficient conversion of cellular motility into persistent tissue-scale migration. These theoretical predictions raised the possibility that suppression of ERK oscillatory coordination may itself contribute to the confinement-induced reduction in migration efficiency. To test this experimentally, we pharmacologically inhibited the ERK pathway using the MEK inhibitor Trametinib in large microstripes. Trametinib treatment induced a rapid decrease in ERK activity and was accompanied by a marked reduction in migration speed toward values closer to those observed under strong confinement **(SI Appendix, Fig. S7)**.

Together, these findings support the idea that confinement suppresses efficient collective migration, at least in part, through progressive damping of mechanochemical ERK oscillatory dynamics. These observations raised the question of how confinement mechanically regulates ERK signaling upstream.

### Cell spreading and pEGFR signaling couple confinement to ERK activity and migration speed

To explore the upstream origin of the confinement–ERK coupling, we examined EGFR activation by immunostaining for phosphorylated EGFR (pEGFR). Representative images revealed strong junctional pEGFR localization in wide stripes **(Fig. 5a–b)**, whereas confinement progressively reduced junctional pEGFR signal intensity **(Fig. 5c–d)**. Recent work has shown that EGFR can be phosphorylated at cell–cell junctions in an E-cadherin–dependent and ligand-independent manner tightly coupled to junction dynamics and cell deformation(19).

**Figure 5.**
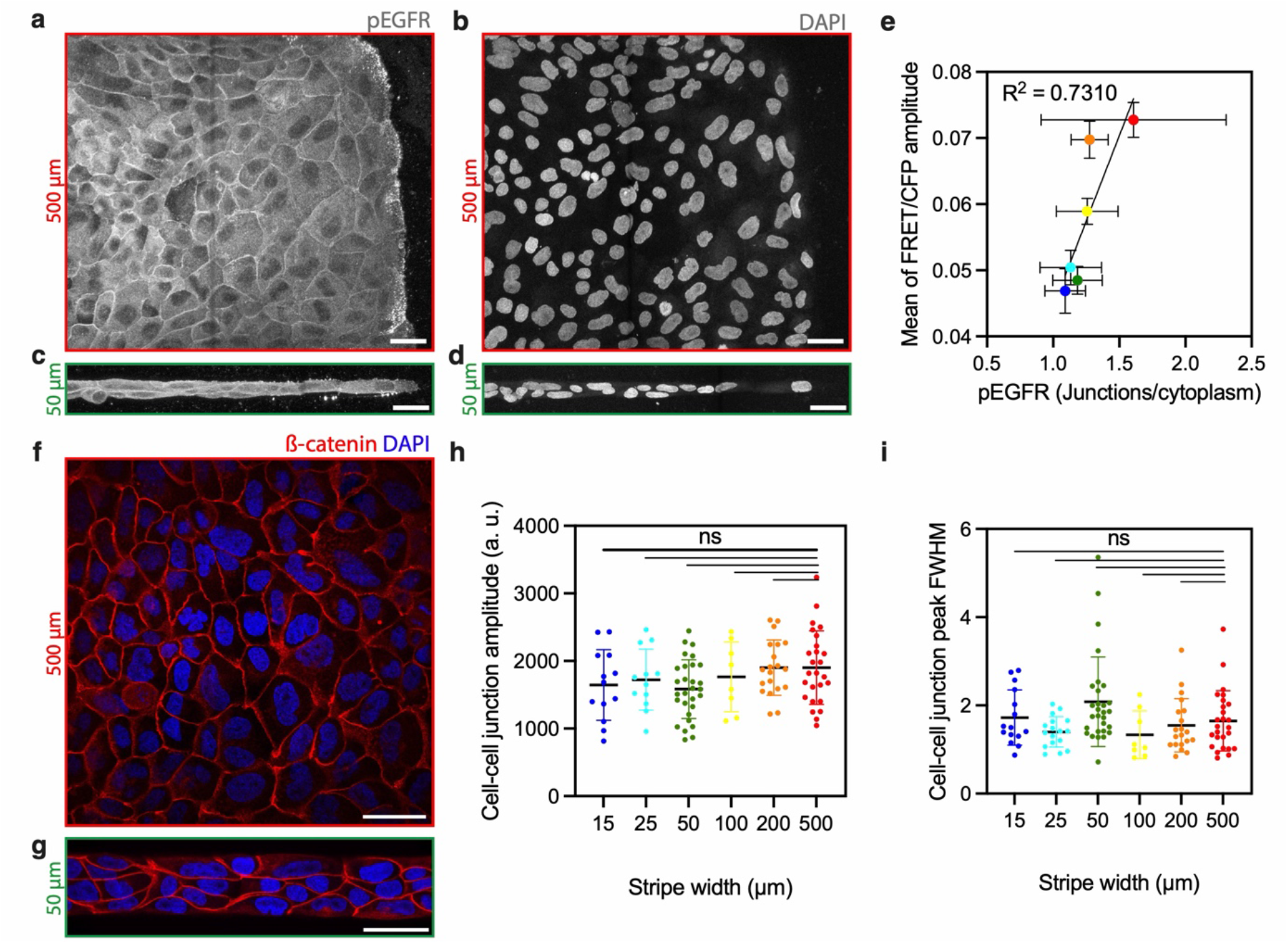
Junctional pEGFR signaling couples confinement to ERK activity. (a–d) MDCK tissues migrating within wide (500 µm; a,b) or confined (50 µm; c,d) microstripes immunostained for pEGFR (a,c) and nuclei (DAPI; b,d), indicating reduced junctional pEGFR enrichment under confinement. Scale bars, 50 µm. (e) Positive correlation between mean ERK activity amplitude and junctional pEGFR levels (junction-to-cytoplasm ratio), revealing a strong association between EGFR activation and ERK signaling (R^2^ = 0.7310). pEGFR measurements were obtained from N=2 replicates, with n = 108 (500 µm), n = 42 (200 µm), n = 29 (100 µm), n = 38 (50 µm), n = 32 (25 µm), and n = 17 (15 µm). (f,g) Representative images of cell–cell junctions visualized by β-catenin staining (red) together with nuclei (DAPI, blue) in MDCK tissues migrating within wide (500 µm; f) or confined (50 µm; g) microstripes. Scale bars, 50 µm. (h) Mean β-catenin junctional fluorescence amplitude as a function of microstripe width. (i) Full width at half maximum (FWHM) of β-catenin junctional fluorescence peaks as a function of microstripe width. Junctional β-catenin measurements were obtained from N=2 replicates, with n = 26 (500 µm), n = 20 (200 µm), n = 8 (100 µm), n = 29 (50 µm), n = 12 (25 µm), and n = 13 (15 µm).

To determine whether this reduction resulted from altered junction organization, we next analyzed cell–cell junction integrity using β-catenin staining **(Fig. 5f-g)**. Quantification of junctional β-catenin signal amplitude and peak width revealed no major changes across confinement conditions **(Fig. 5h-i)**, indicating that the reduction in pEGFR does not arise from junctional disruption itself.

Consistent with this mechanism, reduced cell spreading and junction remodeling under confinement are expected to limit junctional EGFR activation, thereby providing a potential explanation for the decrease in ERK signaling. Quantitative analysis revealed a strong positive correlation between ERK activity amplitude and junctional pEGFR levels **(Fig. 5e**; R^2^ = 0.7310), suggesting that confinement-dependent modulation of EGFR signaling contributes to the regulation of ERK activity.

Together, these results identify cell spreading and EGFR signaling as key intermediates linking spatial confinement to ERK dynamics. We next asked whether this confinement-induced mechanical state is actively controlled by actomyosin contractility and whether it can be reversibly modulated.

### ROCK-dependent contractility modulates the confinement-induced mechanical state and restores ERK signaling and migration

To determine whether the confinement-induced mechanical state is reversible and governed by actomyosin tension, we inhibited ROCK using Y-27632 during collective migration in confined stripes. Live imaging of ERK activity revealed a rapid and pronounced increase in signaling following ROCK inhibition **(SI Movie S8)**, as illustrated by spatial maps along the migration axis **(Fig. 6a)** and representative temporal traces **(Fig. 6b)**. These observations indicate that ERK signaling dynamics remain highly responsive to acute mechanical perturbations even under strong confinement.

**Figure 6.**
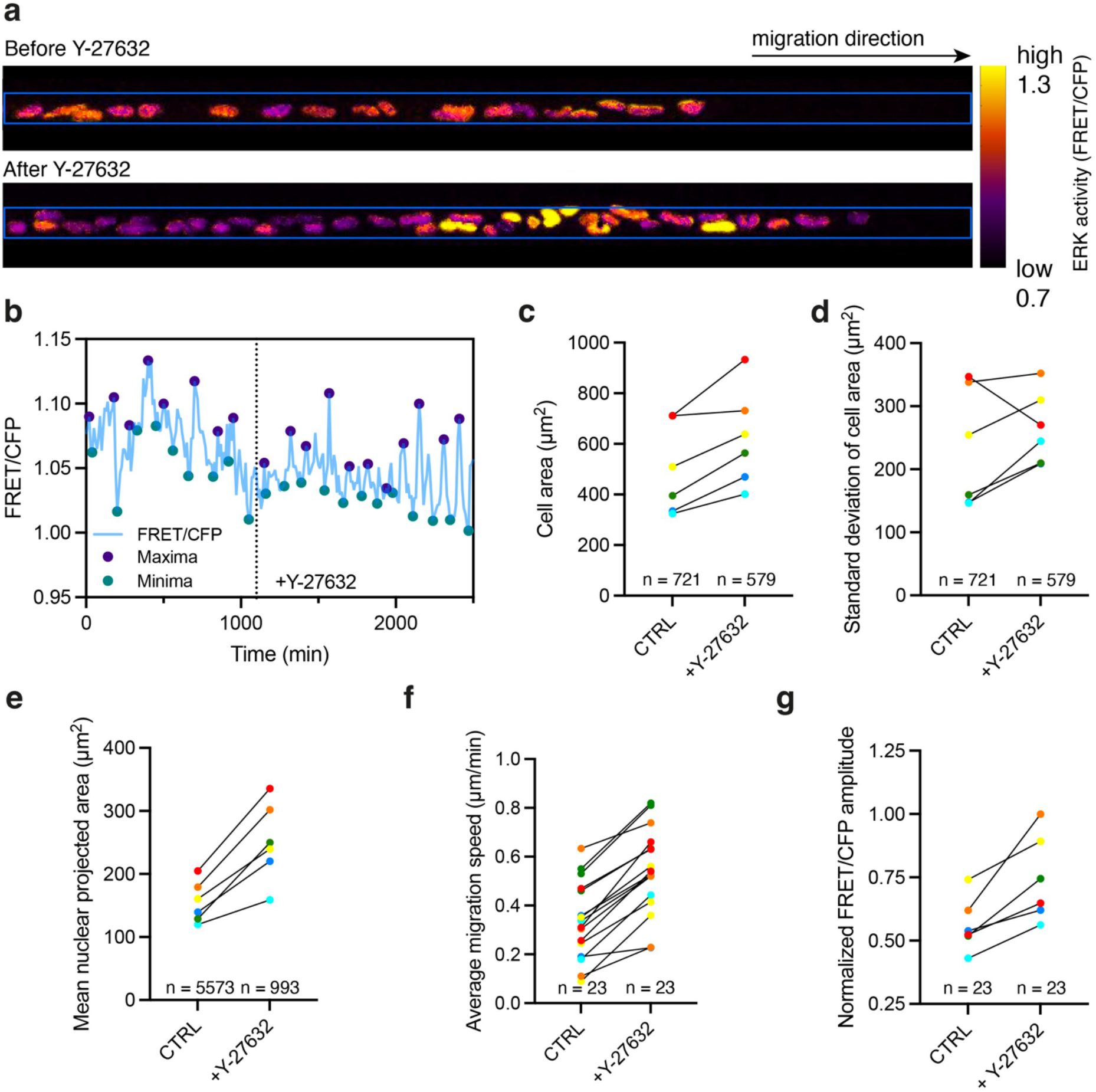
ROCK-dependent contractility modulates the confinement-induced mechanical state and restores ERK signaling and migration. **(a)** Spatial maps of ERK activity (FRET/CFP) along a confined migrating monolayer before and after addition of the ROCK inhibitor Y-27632, illustrating the rapid increase in ERK activity following ROCK inhibition. **(b)** Representative temporal trace of ERK activity before and after Y-27632 addition (dashed line), illustrating rapid enhancement of ERK oscillatory activity during migration in severe confinement conditions (15 µm-wide microstripe). **(c)** Mean cell area and **(d)** standard deviation of cell area in control (CTRL) and Y-27632-treated conditions (+Y-27632), showing increased cell spreading and broader cell area distributions upon ROCK inhibition across confinement conditions. **(e)** Mean nuclear projected area, showing increased nuclear spreading following ROCK inhibition (+Y-27632). **(f)** Average migration velocity and **(g)** normalized FRET/CFP amplitude in control (CTRL) and Y-27632-treated conditions, revealing strong recovery of both ERK signaling and collective migration under confinement following ROCK inhibition (+Y-27632). N ≥ 3 replicates for all conditions.

At the cellular level, ROCK inhibition significantly increased cell spreading across all confinement conditions **(Fig. 6c)**, accompanied by a broader distribution of cell areas **(Fig. 6d)**, consistent with partial release of confinement-imposed mechanical constraints. Consistent with effective inhibition of actomyosin tension, ROCK inhibition markedly reduced phosphorylated myosin light chain (pMLC) levels **(SI Appendix, Fig. S8a**,**b)**. Importantly, this was associated with a marked increase in nuclear projected area **(Fig. 6e)**, indicating that the confinement-imposed mechanical state identified in Fig. 5 is not fixed but is actively tuned by actomyosin contractility.

This was further accompanied by changes in nuclear morphology in three dimensions **(SI Appendix, Fig. S8c**,**d)** and a reduction in nuclear aspect ratio under confinement **(SI Appendix, Fig. S8e)**, supporting a mechanical origin of nuclear deformation. Functionally, these changes resulted in a substantial increase in migration velocity **(Fig. 6f)**, together with a strong recovery of ERK activity amplitude **(Fig. 6g)**, restoring both signaling and collective migration even under highly confined conditions.

Conversely, pharmacological inhibition of the ERK pathway using the MEK inhibitor Trametinib induced a rapid reduction in ERK activity amplitude together with a marked decrease in migration **(SI Appendix, Fig. S7)**, confirming that ERK signaling is required to sustain efficient collective migration.

Together, these results demonstrate that the confinement-induced mechanical state is reversible and governed by ROCK-dependent contractility, and establish that geometric confinement controls collective migration through a contractility-dependent mechanochemical state governing ERK signaling dynamics.

## Discussion

Collective cell migration occurs in geometrically constrained environments where physical boundaries shape tissue organization and dynamics. Here, we show that geometric confinement acts as a key regulator of collective migration by modulating ERK signaling activity. Using controlled micropatterned environments, we demonstrate that increasing confinement progressively reduces ERK activity amplitude and migration efficiency, while preserving the temporal dynamics of ERK signaling. These findings identify ERK activity amplitude as a quantitative readout of the confinement-imposed migratory state.

Our results extend previous work showing that physical confinement regulates epithelial migration by altering cell morphology, adhesion, and force generation (3, 4, 28, 29). While these studies established confinement as a key physical parameter controlling migration efficiency, the link between geometric constraints and intracellular signaling remained unclear. Here, we bridge this gap by demonstrating that ERK signaling is quantitatively modulated by confinement, providing a direct connection between tissue geometry and biochemical signaling during collective migration.

Mechanistically, our data suggest that this coupling emerges from confinement-induced changes in cell spreading and mechanical state. We show that reduced cell spreading is associated with decreased ERK activity, consistent with previous studies linking ERK signaling to cell deformation and mechanical inputs (11, 17). In addition, we find that junctional EGFR activation decreases under confinement, in line with recent work demonstrating that EGFR phosphorylation can occur at cell–cell junctions in a deformation-dependent and ligand-independent manner (19). Together, these observations support a model in which geometric confinement limits cell deformation and junctional dynamics, thereby reducing EGFR activation and downstream ERK signaling.

A key finding of this study is that confinement-induced changes in cell and nuclear geometry provide robust quantitative proxies for the mechanical state imposed by spatial confinement. Despite large changes in nuclear shape and projected area, nuclear volume remains largely constant, indicating that confinement primarily induces anisotropic deformation while preserving nuclear volume. The strong correlations between ERK activity, cell spreading, and nuclear projected area further suggest that epithelial geometry integrates multiple aspects of the mechanical environment and can serve as a quantitative readout of the mechanochemical state of the tissue. This is consistent with growing evidence that the nucleus acts as a central mechanosensor integrating cytoskeletal forces and geometric constraints (30).

ERK signaling is thought to regulate actomyosin contractility in a spatially heterogeneous manner, generating local gradients of pMLC across the tissue (13). Here, we show that the confinement-induced mechanical state is reversible and controlled by actomyosin contractility. Inhibition of ROCK rapidly increases cell spreading, nuclear projected area, ERK activity, and migration velocity, demonstrating that this state is not fixed but dynamically regulated. Conversely, inhibition of ERK signaling reduces migration efficiency, establishing a functional link between ERK activity and collective migration. Together, these results support a model in which geometric confinement regulates collective migration through a contractility-dependent mechanochemical feedback between actomyosin tension and ERK signaling dynamics. Our results therefore suggest that confinement does not merely reduce ERK activity levels but progressively disrupts the collective spatiotemporal organization of ERK waves required for efficient coordinated migration.

More broadly, our findings highlight how physical constraints imposed by tissue architecture can be encoded into intracellular signaling dynamics to regulate collective behavior. In physiological contexts such as development, wound healing, and tumor invasion, cells frequently migrate within confined environments where geometry and mechanics are tightly coupled (8). The identification of a quantitative relationship between confinement, mechanical state, and ERK signaling suggests that epithelial tissues may use such mechanochemical coupling to adapt their migratory behavior to spatial constraints.

Several questions remain for future work. Although our data support a strong association between junctional EGFR activation and ERK dynamics under confinement, additional experiments will be required to determine the precise causal contribution of EGFR signaling to confinement-dependent ERK regulation and to identify how other mechanosensitive pathways may cooperate with EGFR signaling in this context. In particular, it will be important to determine how this mechanochemical coupling operates in three-dimensional environments, where confinement, matrix architecture, and viscoelasticity (31) interact in more complex ways. Extending this framework to pathological contexts such as cancer invasion may also provide important insights into how tumor cells adapt their migration strategies within confined microenvironments.

## Materials and Method

### Patterned substrate preparation

Glass-bottom dishes (IBL) and large Petri dishes were spin-coated (32) with polydimethylsiloxane (PDMS; Sylgard 184, Dow Corning) to generate PDMS-coated substrates and free-standing PDMS sheets, respectively. The PDMS was cured for 6 h at 60 °C. Hemispherical membranes were then cut from the cured PDMS sheets. PDMS stamps containing a large reservoir connected to long stripes of defined widths (500, 200, 100, 50, 25, and 15 µm) were activated by UV/ozone treatment and incubated for 1 h with an aqueous solution of human plasma fibronectin (30 µg/mL; Merck) supplemented with Alexa Fluor 546–conjugated fibrinogen (2 µg/mL; Thermo Scientific). After incubation, the stamps were dried under a nitrogen stream and brought into conformal contact with UV/ozone–activated PDMS-coated dishes to transfer the protein pattern. Following stamp removal, both patterned dishes and PDMS sheets were incubated for 5 min in a 1% (w/w) Pluronic solution (BASF) to prevent nonspecific protein adsorption. The substrates were then thoroughly rinsed with water and dried. Finally, the PDMS sheets were carefully positioned onto the patterned substrates and aligned with the fibronectin stripes.

### Cell culture

Madin–Darby canine kidney (MDCK) cells expressing the EKARrEV-NLS biosensor were maintained in Dulbecco’s modified Eagle’s medium (DMEM; Westburg) supplemented with 10% fetal bovine serum (FBS; Gibco) and 1% penicillin–streptomycin (Westburg), and cultured under standard conditions (37°C, 5% CO_2_). For time-lapse experiments, cells were seeded into the adhesive reservoir at a density of 1.2.10^6^ cells/mL in imaging medium (FluoroBrite DMEM; Gibco) supplemented with FBS and antibiotics. After 24 h, the epithelial tissue formed a cohesive monolayer fully covering the reservoir. The PDMS sheet overlaying the fibronectin stripes was then carefully removed to expose the adhesive patterns. Cells and substrates were gently rinsed several times with phosphate-buffered saline (PBS), and the dish was subsequently filled with fresh imaging medium prior to acquisition.

### Cell staining

MDCK cells were fixed and permeabilized by incubation for 10 min in 4% paraformaldehyde (PFA) containing Triton X-100 (1:2000). Cells were then blocked for 30 min at room temperature in a solution of 5% fetal bovine serum (FBS; Gibco) and 1% bovine serum albumin (BSA; Merck) in phosphate-buffered saline (PBS). Nuclei were stained with 4′,6-diamidino-2-phenylindole dihydrochloride (DAPI; Thermo Scientific) for 45 min. Cell–cell junctions were labeled using a mouse anti–β-catenin primary antibody (Santa Cruz; 1:200, 45 min incubation), followed by an Alexa Fluor 555–conjugated goat anti-mouse IgG (H+L) secondary antibody (Thermo Scientific; 1:200, 45 min incubation). Phosphorylated EGFR (pEGFR) was detected using a rabbit anti–EGFR (phospho-Y1068) antibody (Abcam; 1:200, 45 min incubation), followed by an Alexa Fluor 647–conjugated goat anti-rabbit IgG (H+L) secondary antibody (Abcam; 1:200, 45 min incubation). Phosphorylated myosin light chain (pMLC) was detected using a rabbit anti–phospho-myosin light chain 2 (Thr18/Ser19) antibody (Cell Signaling Technology; 1:200, 45 min incubation), followed by an Alexa Fluor 647–conjugated goat anti-rabbit IgG (H+L) secondary antibody (Abcam; 1:200, 45 min incubation).

### Drug treatments

The inhibition of MEK (Mitogen-Activated Protein Kinase 1) was performed with a treatment with Trametinib (GSK1120212, Selleck Chemicals) added to the cell culture medium for a final concentration of 100 nM. The inhibition of ROCK was performed with a treatment with Y-27632 (Sigma-Aldrich) added to the cell culture medium for a final concentration up to 50 µM.

### Time-lapse imaging and confocal microscopy

MDCK cell migration within confinement stripes was imaged at ×10 or ×20 magnification using a Photometrics Prime 95B camera (Photometrics, Tucson, AZ) mounted on a motorized inverted microscope (Nikon Ti2 equipped with an A1R HD25 system). An environmental control chamber (Okolab) was used to maintain cells at 37 °C and 5% CO_2_ throughout imaging. Time-lapse images were acquired over extended periods (24 to 100 h) in both differential interference contrast (DIC) and fluorescence modes, with a time interval of 10 min between frames. FRET imaging was performed using a dedicated filter set (CFP-2432C, 32 mm). The donor fluorophore (CFP) was excited using a 438/24 nm BrightLine bandpass filter in combination with a 458 nm BrightLine single-edge dichroic mirror. Emission signals from both the donor (CFP) and acceptor (YFP) were acquired sequentially, with acceptor emission collected through a 542/27 nm BrightLine bandpass filter. Acquisition parameters were kept constant across all conditions, and ERK activity was quantified as the ratio of acceptor to donor fluorescence (YFP/CFP). Three-dimensional confocal imaging was performed on a Nikon Ti2 A1R HD25 confocal microscope using Plan Apochromat silicone immersion objectives: ×40 (NA = 1.25, WD = 0.30 mm) and ×100 (NA = 1.35, WD = 0.30 mm). Z-stacks were acquired with a step size of 0.125 µm using NIS-Elements software (Advanced Research v6.1).

### Image and data analysis

The ratio of FRET to CFP fluorescence (FRET/CFP) was quantified within regions of interest (ROIs) containing a small number of cells, positioned at the center of each confinement stripe approximately 500 µm from the stripe entrance. FRET/CFP ratios were computed using a custom MATLAB (MathWorks) script adapted from previously published work (33). In the corresponding maps, color encodes the FRET/CFP ratio. Temporal variations in FRET/CFP signals were analyzed using a custom Python pipeline based on the pandas, NumPy, SciPy, and matplotlib libraries. Time-series data were imported from Excel files containing fluorescence intensity measurements. Local maxima and minima were identified using the find peaks function from the SciPy library, with a minimum peak distance constraint defined by the temporal resolution to exclude noise-related fluctuations. Maxima and minima were paired based on temporal proximity, and ERK activity amplitude was calculated as the difference between each maximum and its nearest minimum. The time interval between successive maxima was used to estimate oscillation periodicity. Extracted parameters were compiled and exported for further analysis. Data visualization was performed using matplotlib.

Cell segmentation for cell area heatmaps was performed using MorphoGraphX. Nuclear segmentation for quantification of nuclear area and aspect ratio was carried out using FIJI (ImageJ) with Cellpose-based segmentation applied to epifluorescence images or time-lapse sequences. The cell shape index (CSI) was calculated from the following relationship: CSI = 4 π×area/(perimeter)^2^. The CSI ranges from 1 for a perfectly circular shape to 0 for a highly elongated morphology. Nuclear volumes were quantified from 3D confocal stacks of DAPI-stained samples using the 3D object measurement plugin in NIS-Elements (AR v6.1). Velocity vector fields of tissues confined in stripes were computed using PIVlab (MATLAB, MathWorks). Time-lapse fluorescence images of cell nuclei were analyzed using a two-pass fast Fourier transform (FFT) window deformation algorithm. Interrogation windows of 76×76 µm and 38×38 µm were used for the first and second passes, respectively, with 50% overlap. Displacement fields were converted into physical units using spatial calibration, and velocity magnitude and direction were extracted from the resulting vector fields. For pEGFR quantification, fluorescence intensity profiles were extracted using line plot profile analysis across cell–cell junctions and adjacent cytoplasmic regions after subtraction of the substrate background signal. Junction-to-cytoplasm ratios were then calculated by normalizing the mean junctional pEGFR intensity to the corresponding cytoplasmic intensity measured in the same region.

### Statistical analysis

Statistical significance was assessed using one-way analysis of variance (ANOVA) followed by Tukey’s post hoc test for multiple comparisons. Significance levels were defined as P < 0.05 (*), P < 0.01 (**), P < 0.001 (***), and P < 0.0001 (****). Non-significant differences are not indicated. Unless otherwise stated, data are presented as mean ± standard deviation (s.d.).

## Acknowledgments

M.V. and S.G. acknowledge funding from the University of Mons–UMONS, the F.R.S.-FNRS Epiforce Project no. T.0092.21, the F.R.S.-FNRS Cellsqueezer project no. J.0061.23, and the F.R.S.-FNRS Optopattern project no. U.NO26.22. This work was supported by the Interreg projects ANTIRESI and MICROPLAITE, cofunded by the European Union through the European Regional Development Fund (ERDF-FEDER) under the Interreg France-Wallonie-Vlaanderen Program. M.V and S.G. acknowledge the SPW Research Win^2^WAL Program INERVODERM, funded by the Walloon Region. S.G. acknowledges le Fonds pour la Recherche Médicale dans le Hainaut (FRMH) and support from the Francqui Foundation as a Francqui Research Professor. T.H., M.L. and S.G. thank ASEM-DUO Fellowship programme “DUO Wallonia-Brussels” (Belgium).

## Author contributions

S.G. and M.V. conceived the project, and S.G. supervised the study. M.V. developed the micropatterning strategy and performed the time-lapse migration experiments, cell-tracking analyses, immunostaining, and confocal imaging. S.G. and M.V. analyzed and interpreted the data and prepared the figures. E.H. developed the theoretical framework and performed the simulations. R.T. contributed to data analysis and provided critical feedback on the manuscript. M.L. initiated the collaboration with T.H. on ERK signaling and provided critical feedback on the manuscript. T.H. provided the ERK-MDCK cell line, contributed to the optimization of imaging conditions, and developed the MATLAB algorithms used for ERK signal analysis. The manuscript was written by M.V. and S.G., with input from all authors. All authors contributed to data interpretation, revised the manuscript, and approved the final version.

## Data and materials availability

All data needed to evaluate the conclusions in the paper are present in the paper and/or the materials cited herein.

