## Supplementary Information for "Geometric confinement regulates ERK signaling dynamics and collective migration"

### 1. Supplementary Theory

Here, we use a minimal extension of the mechanochemical wave model of Boocock *et al.* (Nat. Phys. 2021) to ask whether the experimentally observed loss of ERK oscillations under lateral confinement can be accounted for by the published model under the imposed change in mean cell area. We find that the same three coupled equations that produce ERK waves in unconfined epithelia can indeed predict that uniform compression shifts the operating point of the dynamical system across a Hopf bifurcation, suppressing oscillations while leaving their intrinsic period largely unchanged. No additional ingredients beyond the original Boocock framework are required. This is the same phenomenology as the well-known suppression of repetitive firing in FitzHugh–Nagumo and related excitable models when the input bias is moved outside an oscillatory window.

The equations for ERK mechanochemical waves read, at the continuum level,

$$\begin{cases} \tau_r \partial_t r = \partial_{xx} r - \partial_x l_0 \\ \tau_l \partial_t l_0 = -(l_0 - 1) - \alpha(ERK - ERK_0) \\ \tau_E \partial_t ERK = -(ERK - ERK_0) - (ERK - ERK_0)^3 + \beta \partial_x r \end{cases}$$

where ERK is the local ERK activity, $l_0$ the prefered diameter of a cell and $r$ the position of a cell (so that the actual diameter of a cell is $\partial_x r$. As previously shown, this model admits spatio-temporal waves with finite wavelength and period above a critical value of $\alpha\beta$, at which the system undergoes a supercritical Hopf bifurcation. This can be seen by linearizing the system around its prefered steady state values of $ERK = ERK_0$, $l_0 = 1$. Following Boocock et al, we integrated the discrete chain equations using an explicit Euler scheme (time step $dt$ = 0.1min, $N$ = 100 cells, stress-free boundary conditions), with parameters in the supercritical regime at zero compression: $\tau_r$ = 2min, $\tau_\ell$ = 40min, $\tau_E$ = 5min, $\alpha$ = 15, $\beta$ = 3, $ERK_0$ = 1 [all taken from (17)]. Random initial conditions were equilibrated at for $T_{eq}$ = 300 min. To understand the effect of confinement, we then applied global compression to the system (effectively restricting the total area accessible to cells). We did so very slowly so that dynamics are quasi-static, i.e. we can assess the behavior of ERK waves for every compression value. These simulations reproduce the experimental observations of (i) confinement-dependent reduction of ERK oscillation amplitude, (ii) preservation of intrinsic period of the ERK waves, and (iii) abrupt loss of oscillations beyond a critical confinement. Within this framework, the rescue of ERK signalling by ROCK inhibition is naturally interpreted as a release of the imposed mean strain. Pharmacological reduction of contractility allows cells to respread, and the system re-enters the supercritical Hopf regime of mechanochemical ERK waves. This can also be understood analytically by a simple linear stability analysis of our continuum equations. If confinement is exerted on the system, by a strain $\epsilon_c$, then the system reaches a different steady state $ERK_0 - ERK^*$, which satisfies $ERK^* + {ERK^*}^3 = \beta\epsilon_c$ and $l_0 = 1 - \alpha\, ERK^*$. However, linearizing around this regime stiffens the effective relaxation rate of ERK, by a factor $1 + 3\, {ERK^*}^2$, thus dampening the ERK waves and increasing the effective threshold of the instability. This generically predicts that confinement will move the system away from the region of spontaneous pattern formation.

Next, we asked how the suppression of waves connects to migration efficiency. Previously (17), it was also shown in confined colonies that the polarization field underlying active cell migration p could be described by the following equation:

$$\tau_p \partial_t p = -p + \gamma(ERK)\partial_x \sigma_{xx}$$

where polarity relaxes to a zero value with time scale $\tau_p$ in the absence of any external signal, but is trigerred by gradients of mechanical stress (directionality) and modulated by local ERK levels (amplitude, with $\gamma(ERK)$ being a monotonouly decreasing function). ). In the presence of directional ERK waves, directional stress waves $\sigma_{xx}$ are rectified by $\gamma(ERK)$, which gives rise to directional motion. Thus, in these equations, if confinement kills ERK waves, the $\sigma_{xx}$ term becomes close to spatially constant, so that $\partial_x \sigma_{xx} \approx 0$. In this case, the total average of polarized traction force $p$ goes to 0, even if absolute local values of traction forces (represented by $\gamma$) would go up. Given that overall tissue velocity scales with the magnitude of polarized traction forces $p$, this can explain how the loss of directional ERK waves triggered by confinement leads to a drastic slow-down of collective cell migration in thin microstripes.

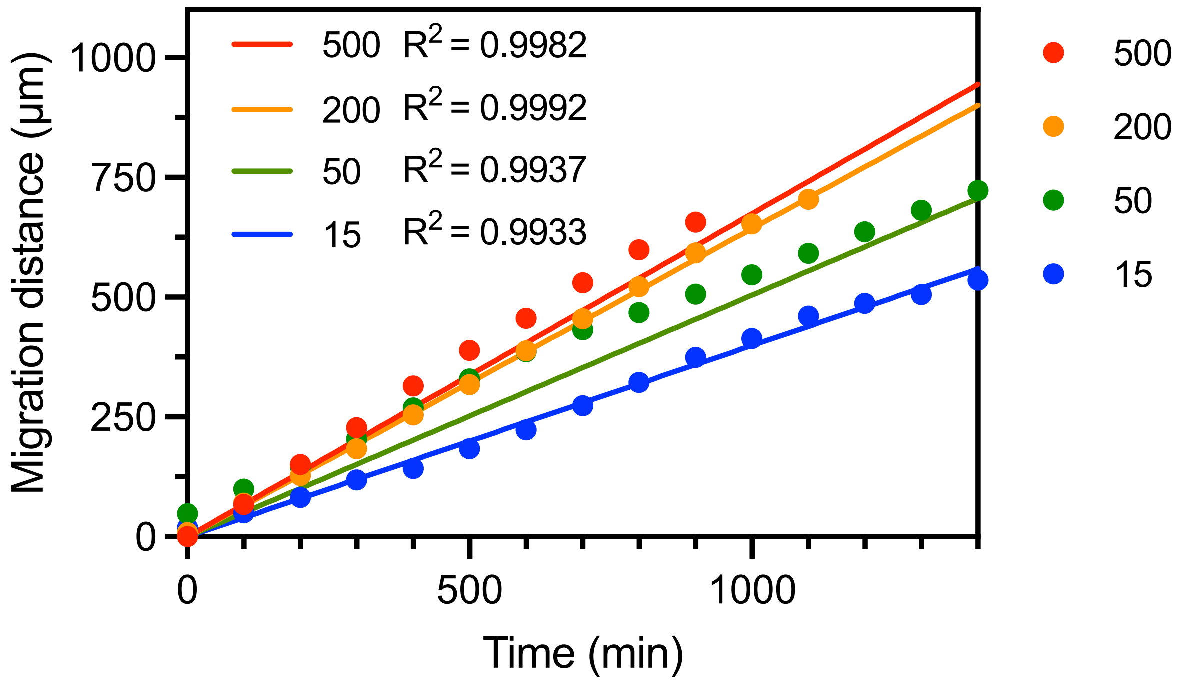


**Supplementary Figure S1 – Temporal evolution of collective migration under different confinement conditions.** Migration distance as a function of time for epithelial MDCK monolayers entering adhesive microstripes of widths 500 µm (red), 200 µm (orange), 50 µm (green), and 15 µm (blue). Solid lines represent linear regressions of the mean ± SD obtained from multiple independent experiments, revealing a progressive reduction in migration speed with increasing confinement. Data were obtained from n = 9 microstripes (500 µm), n = 13 (200 µm), n = 10 (50 µm), and n = 5 (15 µm), with N ≥ 3 replicates for all conditions.

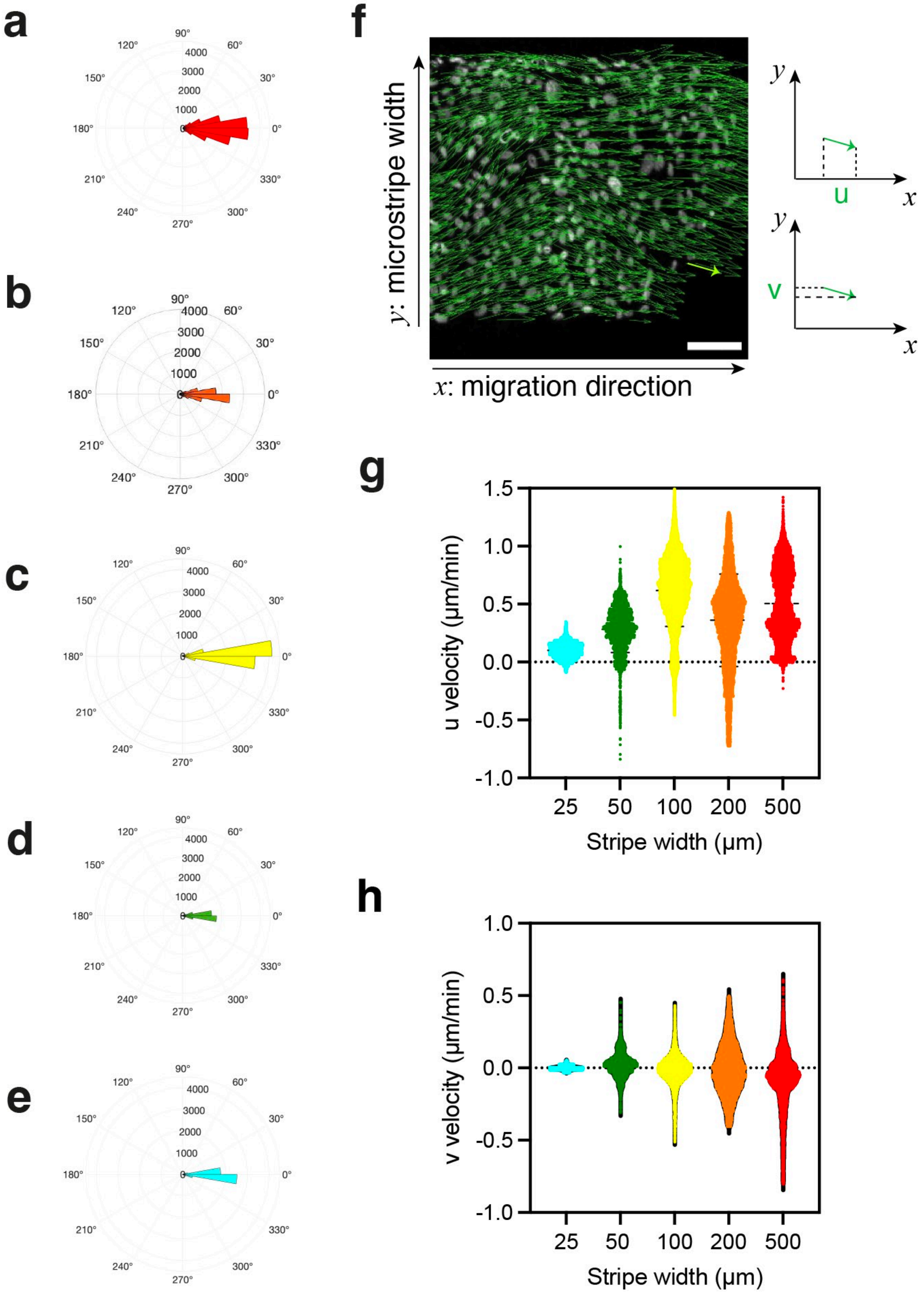


**Supplementary Figure S2 – PIV analysis reveals confinement-dependent alignment of collective migration flows. (a–e)** Angular distribution of PIV velocity vectors relative to the long axis of the micros for MDCK tissues migrating within microstripes of varying widths: 500 μm (red, a), 200 μm (orange, b), 100 μm (yellow, c), 50 μm (green, d), and 25 μm (turquoise, e). Increasing confinement progressively aligns migration flows along the microstripe axis. **(f)** Schematic representation of the longitudinal (u) and transverse (v) velocity components extracted from PIV analysis relative to the migration direction. **(g)** Distribution of the longitudinal velocity component (u) along the microstripe axis for MDCK tissues migrating within microstripes of varying widths. **(h)** Distribution of the transverse velocity component (v) perpendicular to the microstripe axis, showing progressive suppression of lateral migration flows under strong confinement. u and v velocity components were extracted from a single time-lapse experiment, with $n = 9.985\times10^3$ (500 μm), $n = 14.082\times10^3$ (200 μm), $n = 15.281\times10^3$ (100 μm), $n = 3.491\times10^3$ (50 μm), and $n = 15.976\times10^3$ (25 μm).

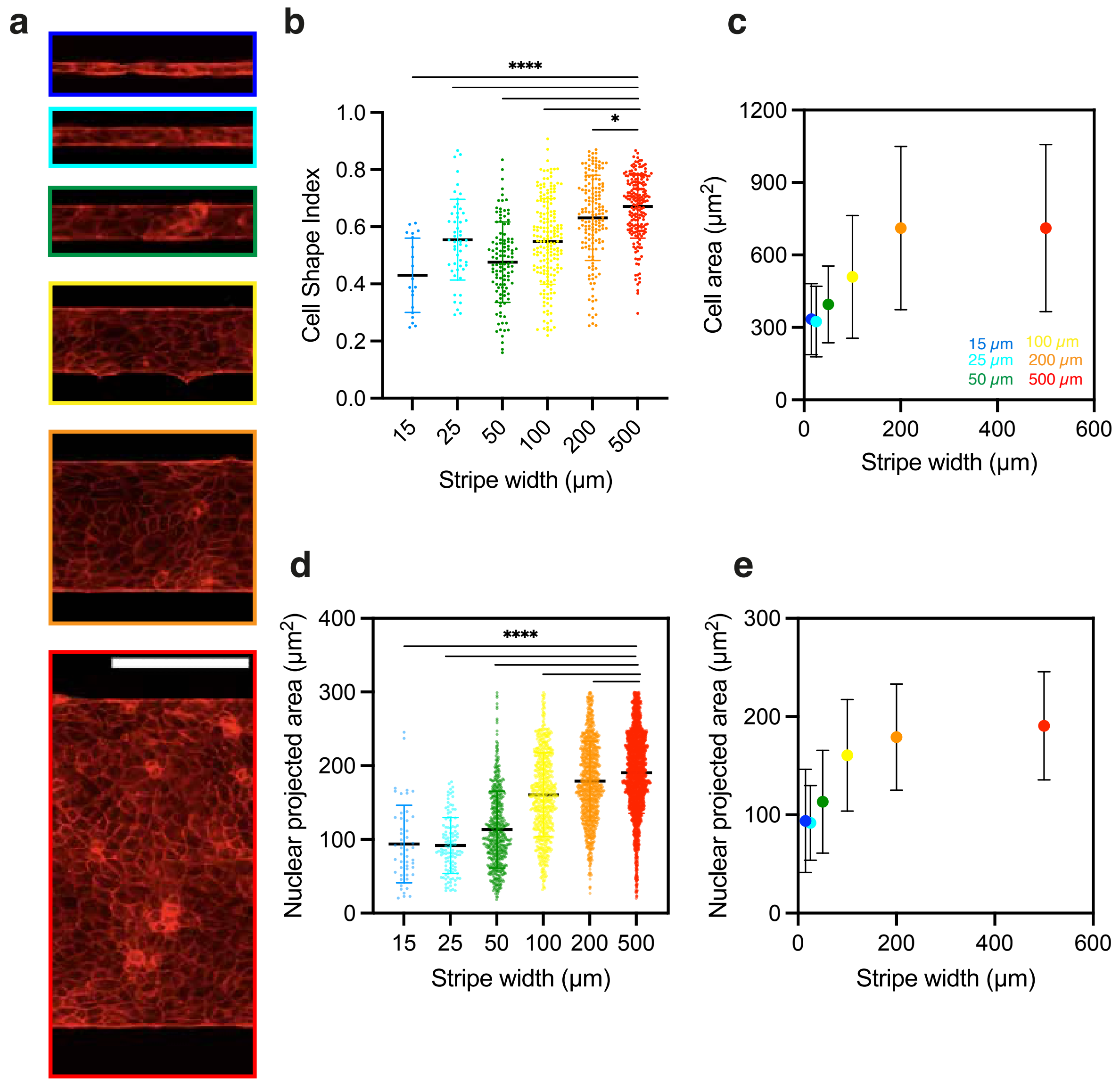


**Supplementary Figure S3 – Geometric confinement progressively reshapes cellular and nuclear morphology**. **(a)** Typical epifluorescence images of MDCK tissues migrating within microstripes of varying widths: 15 µm (blue), 25 µm (turquoise), 50 µm (green), 100 µm (yellow), 200 µm (orange), and 500 µm (red), visualized by β-catenin staining of cell-cell junctions. Scale bar, 200 µm. **(b)** Cell shape index and **(c)** mean cell area as a function of microstripe width during confined migration. **(d)** Distribution of nuclear projected area across microstripe widths, showing progressive reduction in nuclear projected area under confinement. **(e)** Mean nuclear projected area as a function of microstripe width. Cell morphology measurements were obtained from N=3 replicates, with n= 197 (500 µm), n= 155 (200 µm), n= 186 (100 µm), n= 111 (50 µm), n= 52 (25 µm) and n= 20 (15 µm).

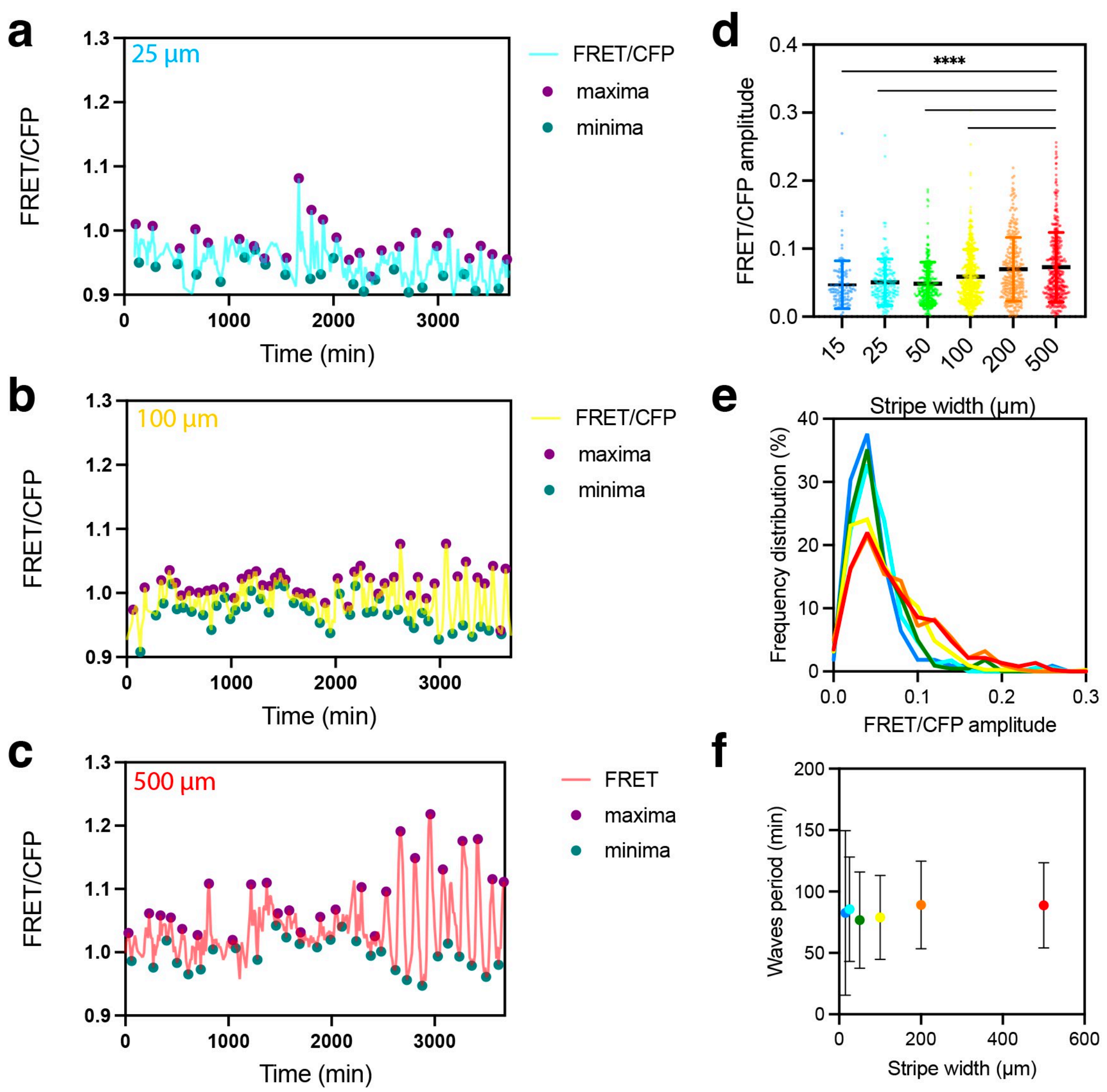


**Supplementary Figure S4 – Confinement progressively dampens ERK oscillatory activity without altering wave period. (a–c)** Temporal evolution of the FRET/CFP ratio measured within the ROI described in Fig. 4b during collective migration in microstripes of (a) 25 µm, (b) 100 µm, and (c) 500 µm widths. Local maxima (purple) and minima (green) were identified using a custom Python script to quantify ERK oscillation amplitudes. **(d)** Mean FRET/CFP oscillation amplitude as a function of microstripe width in collectively migrating MDCK cells, showing progressive damping of ERK oscillations under confinement. **(e)** Frequency distribution of ERK activity amplitudes across microstripe widths, revealing a progressive shift toward lower oscillation amplitudes with increasing confinement. **(f)** Time interval between successive ERK activity maxima (defined as the oscillation period) as a function of microstripe width, revealing that ERK oscillation periodicity remains largely independent of confinement. FRET/CFP signals were acquired from N replicates, with a total of n measurements: N = 9 and n = 373 (500 µm), N = 7 and n = 279 (200 µm), N = 9 and n = 411 (100 µm), N = 5 and n = 222 (50 µm), N = 6 and n = 171 (25 µm), and N = 4 and n = 109 (15 µm).

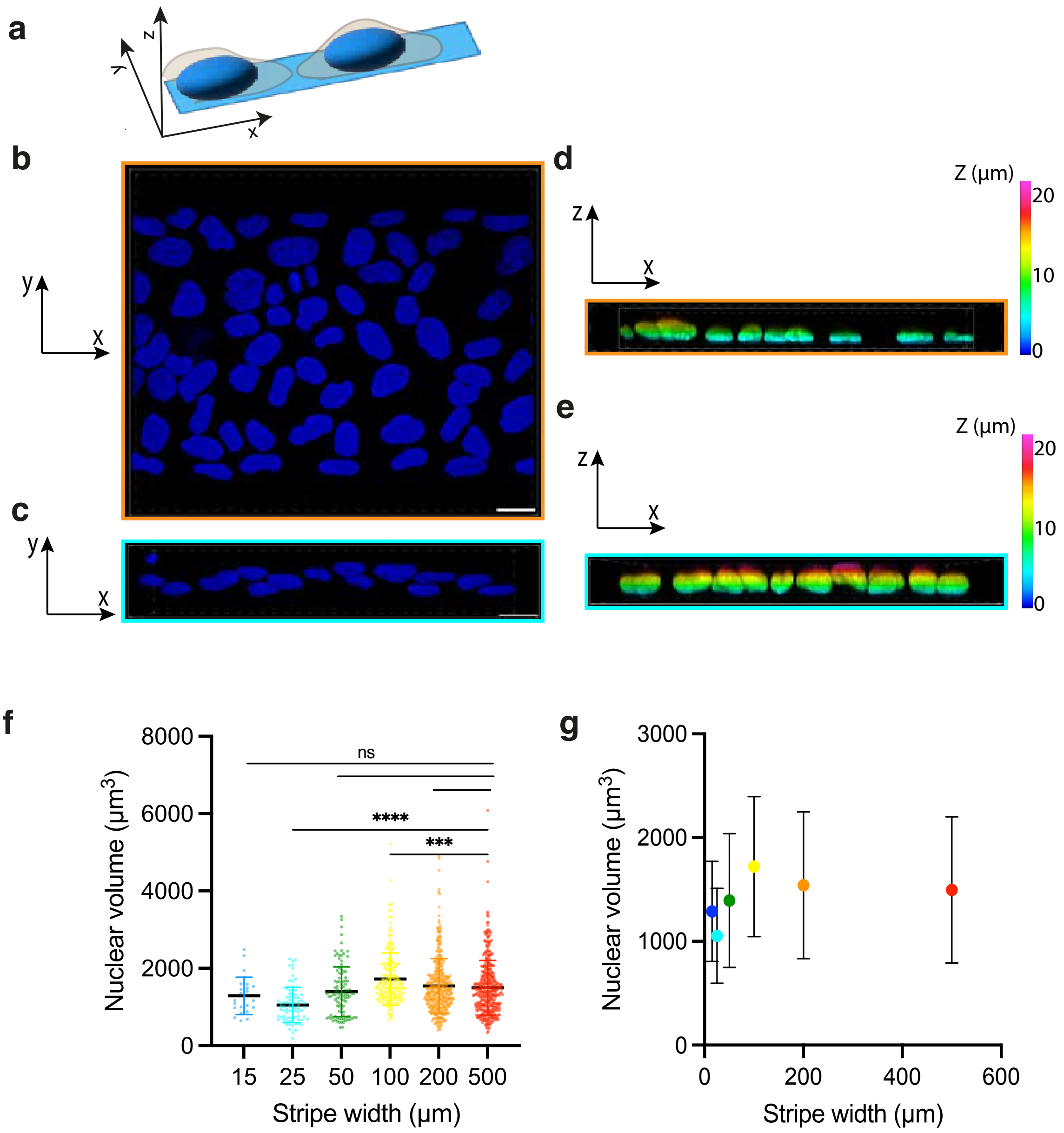


**Supplementary Figure S5 – A reduced nuclear projected area imposed by 2D confinement correlates with decreased ERK activity**. **(a)** Schematic representation of the 3D imaging axes. (b, c) XY top views of confocal α-renderings of nuclei (blue, DAPI staining) in MDCK tissues migrating on **(b)** 200 µm or **(c)** 15 µm wide microstripes. Scale bars, 25 µm. **(d, e)** Z-depth color-coded XZ views of confocal α-renderings of nuclei (blue, DAPI staining) in MDCK tissues migrating on **(d)** 200 µm or **(e)** 15 µm wide microstripes. **(f)** Nuclear volume as a function of microstripe width during confined migration. **(g)** Same data as in (f), displayed with the x-axis as a continuous linear scale. Nuclear volume measurements were obtained from n = 427 nuclei (500 µm), n = 372 (200 µm), n = 216 (100 µm), n = 106 (50 µm), n = 78 (25 µm), and n = 23 (15 µm) microstripes.

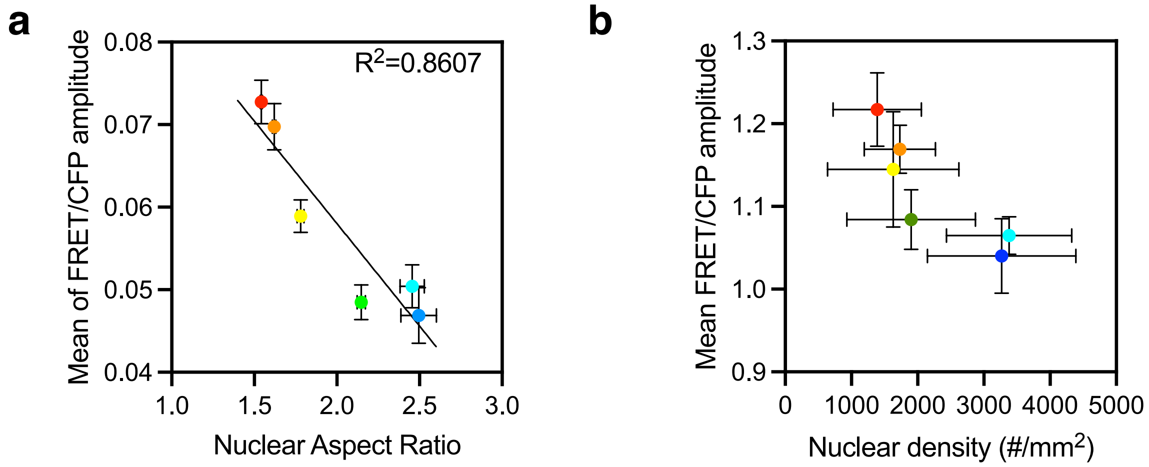


**Supplementary Figure S6 – ERK activity scales with nuclear geometry and crowding imposed by confinement. (a)** Mean ERK activity amplitude (FRET/CFP) as a function of nuclear aspect ratio, revealing a strong positive correlation ($R^2 = 0.8607$), indicating that nuclear deformation provides a geometric proxy for the confinement-imposed mechanical state. **(b)** Mean ERK activity amplitude (FRET/CFP) as a function of nuclear density, revealing a nonlinear dependence on tissue crowding, with ERK activity decreasing primarily at low-density regimes associated with weak confinement.

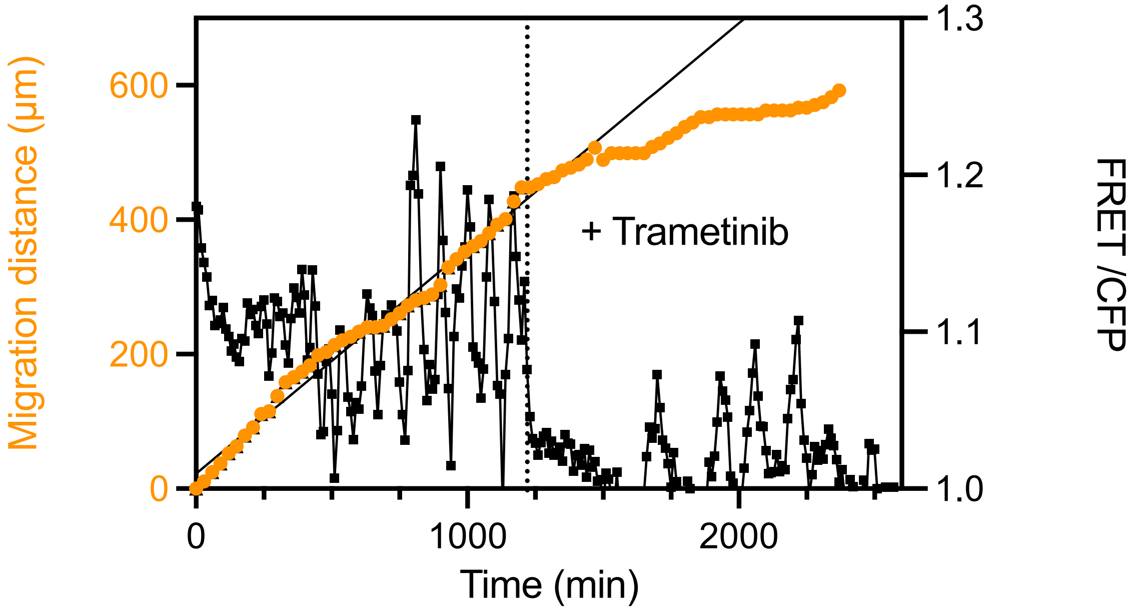


**Supplementary Figure S7 – Trametinib-mediated MEK inhibition suppresses ERK activity and strongly impairs collective migration**. Temporal evolution of migration distance (orange, left axis) and ERK activity (black, right axis; FRET/CFP ratio) measured during collective migration of an MDCK monolayer within a 200 µm-wide microstripe. Trametinib (100 nM) was added after 1,220 min of control migration (dashed line), resulting in a rapid suppression of ERK activity together with a marked reduction in migration progression.

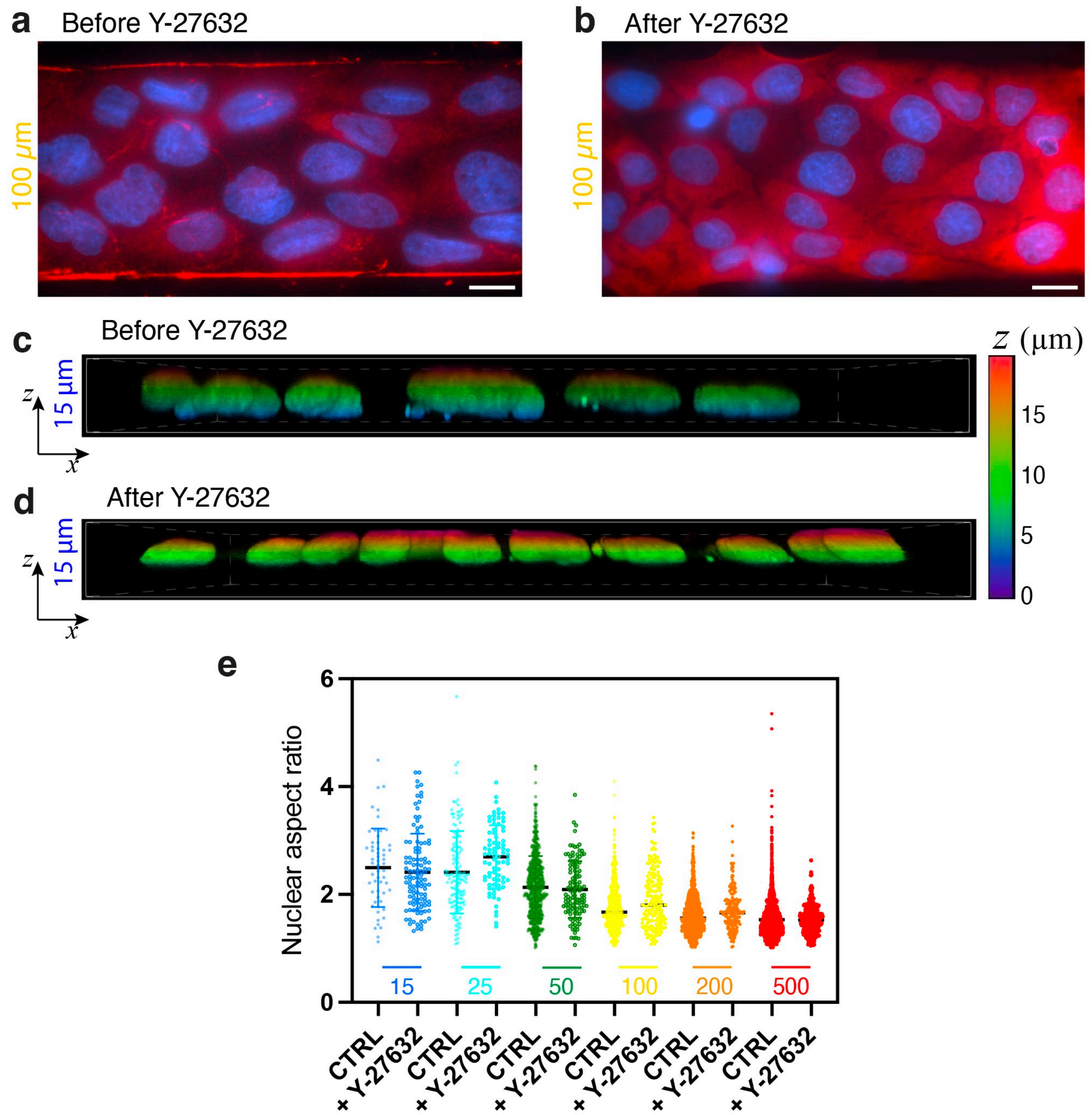


**Supplementary Figure S8 – ROCK inhibition modulates cell and nuclear morphologies under confinement. (a, b)** Representative images of nuclei (DAPI, blue) and phosphorylated myosin light chain (pMLC, red) in MDCK tissues migrating within 100 µm-wide microstripes before (a) and after (b) ROCK inhibition with Y-27632. ROCK inhibition reduces pMLC levels and is associated with increased cell spreading. Scale bars, 20 µm. **(c, d)** Z-depth color-coded XZ views of confocal α-renderings of nuclei (DAPI staining) in MDCK tissues migrating within 15 µm-wide microstripes before (c) and after (d) ROCK inhibition with Y-27632, revealing reduced vertical nuclear confinement following ROCK inhibition. **(e)** Nuclear aspect ratio as a function of microstripe width during confined migration in control (CTRL) and Y-27632-treated tissues, showing conserved nuclear elongation upon ROCK inhibition across confinement conditions.

**Supplementary Movie S1 –** Time-lapse DIC imaging of MDCK monolayers migrating within adhesive microstripes of widths (from top to bottom): 500, 200, 100, 50, 25, and 15 µm. Scale bar, 200 µm. Time is shown as hh:mm:ss.

**Supplementary Movie S2 –** Time-lapse imaging of MDCK nuclei visualized using the EKARrEV-NLS probe during collective migration within a 500 µm-wide microstripe. PIV velocity vectors are overlaid in green. Scale bar, 100 µm. Time is shown as hh:mm:ss.

**Supplementary Movie S3 –** Time-lapse imaging of MDCK nuclei visualized using the EKARrEV-NLS probe during collective migration through a funnel-shaped adhesive micropattern composed of successive regions of decreasing width (500, 200, 100, 50, 25, and 15 µm). Scale bar, 200 µm. Time is shown as hh:mm:ss.

**Supplementary Movie S4 –** Three-dimensional confocal rendering of DAPI-stained nuclei in MDCK cells migrating within a 200 µm-wide microstripe.

**Supplementary Movie S5 –** Three-dimensional confocal rendering of DAPI-stained nuclei in MDCK cells migrating within a 25 µm-wide microstripe.

**Supplementary Movie S6 –** Spatiotemporal modulation of ERK activity in an MDCK monolayer migrating within a 500 µm-wide microstripe. ERK activity is visualized as a color-coded FRET/CFP ratio. The white box indicates the region of interest (ROI) used for quantification of ERK activity. Scale bar, 100 µm. Time is shown as hh:mm:ss.

**Supplementary Movie S7 –** Spatiotemporal modulation of ERK activity in an MDCK monolayer migrating within a 25 µm-wide microstripe. ERK activity is visualized as a color-coded FRET/CFP ratio, revealing dampened oscillatory dynamics under strong confinement. Scale bar, 50 μm. Time is shown as hh:mm:ss.

**Supplementary Movie S8** – Spatiotemporal modulation of ERK activity in an MDCK monolayer migrating within a 15 μm-wide microstripe, visualized as a color-coded FRET/CFP ratio. The ROCK inhibitor Y-27632 was added after 18 h of imaging, resulting in a rapid increase in ERK activity. Scale bar, 100 μm. Time is shown as hh:mm:ss.
